# A small lipidated peptide targeting Na_V_1.8 channels attenuates osteoarthritic pain behavior and prevents bone erosion

**DOI:** 10.1101/2025.01.28.635324

**Authors:** Raider Rodriguez, Joseph B. Kuechle, Arin Bhattacharjee

## Abstract

Chronic pain is a major concern for patients with osteoarthritis (OA). Current treatments for OA-related pain often fail and may worsen the disease. The sodium channel subtype Na_V_1.8 has emerged as a potential target for treating pain, leading to the development of Na_V_1.8 inhibitors in clinical trials. We previously identified Magi-1, a WW domain-containing scaffold protein, as a regulator of Na_V_1.8 at the plasma membrane of nociceptive neurons. Disrupting the interaction between Na_V_1.8 and Magi-1 facilitated channel degradation in neurons, reducing pain behavior in multiple animal models. In this study, we investigated the impact of disrupting Na_V_1.8 scaffolding on an animal model of OA pain using genetic and pharmacological approaches. Genetic Magi-1 knockdown effectively attenuated established OA pain in mice. Pharmacological targeting of the Na_V_1.8-Magi-1 interaction in rats with a lipidated Na_V_1.8 WW binding domain decoy peptide inhibited pain behavior for multiple weeks. MicroCT imaging revealed minimal alterations in subchondral bone remodeling in animals injected with the lipidated decoy peptide compared to those receiving scrambled peptide-control animals. This suggested that the Na_V_1.8 peptidomimetic not only alleviated OA pain but also delayed joint degeneration. Our preclinical studies indicate that intraarticular injection of lipidated peptides capable of disrupting ion channel scaffolding in neurons can provide effective and sustained analgesia for several weeks after a single administration.

## Introduction

Osteoarthritis (OA) is the most prevalent cause of disability among the elderly population, not only posing a public health concern for individuals but also contributing to socioeconomic burdens. Around 595 million people worldwide were living with OA in 2020, a number that is projected to increase significantly by 2050^1,2^. Chronic pain remains the primary symptom among OA patients, which is characterized by inflammatory and neuropathic components at the various stages of the disease^3^. There are multiple analgesic approaches for OA including total joint replacement^4^, but injectable therapies are more attractive to oral medications because of a safer side effect profile and are much less invasive than arthroplasty^5^. However, a randomized clinical trial demonstrated that local injection of the corticosteroid triamcinolone yielded no significant difference in pain scores compared to placebo. Furthermore, triamcinolone was observed to exacerbate joint integrity over the long term^6^.

Approximately 70-80% of the knee joint’s nociceptors are composed of small-diameter unmyelinated C fibers. These fibers preferentially express Na_V_1.8 sodium channels and intra-articular injection of an Na_V_1.8 inhibitor was shown to reduce OA pain behavior^7^. The effects were short-lived^7^ presumably because the drug washes away from the site of injection. Consequently, there has been a concerted effort to develop systemic Na_V_1.8 pore blockers. Recent clinical trials suggest that one potent Na_V_1.8 blocker called suzetrigine, although with a modest effect, significantly reduced acute human pain^8,9^. Interestingly, it was shown that Na_V_1.8 blockers, including suzetrigine, exhibit a property of “reverse use-dependent inhibition”, where inhibition by drug is relieved by depolarization^10,11^. This might explain why the highest dose of drug was necessary to achieve the modest clinical effect for acute pain^8^. A more concerning finding was reported in a Phase II trial involving patients with diabetic neuropathy who received high-dose suzetrigine treatment over a 12-week period: this treatment resulted in a decreased creatinine clearance^12^. We have taken a different approach to target Na_V_1.8 channels in C fibers^13^. We previously identified the WW domain containing scaffold protein Magi-1, as an interactor of Na_V_1.8 channels in dorsal root ganglion (DRG) neurons^14^. Na_V_1.8 channels contain a WW domain binding motif (PPXY) in its C-terminal domain which is predicted to bind to proteins that contain WW domains such as Magi-1 and the E3 ubiquitin ligase Nedd4-2^15^. We observed that *in vivo* Magi-1 knockdown in DRG neurons resulted in a decrease in Na_V_1.8 expression and reduction in pain behavior^14^. This finding suggested that Magi-1 and E3 ubiquitin ligases compete for binding to Na_V_1.8 channels and Magi-1 functions as a protective factor for Na_V_1.8 channels against ubiquitin-mediated degradation^16^. Subsequently, we engineered a small, lipidated decoy peptide derived from Na_V_1.8 PPXY motif to locally disrupt Magi-1 and Na_V_1.8 interaction^14^ to facilitate the loss of Na_V_1.8 channels in DRG neurons^13^. Lipidation of this Na_V_1.8 targeted peptide enables neuronal membrane penetration and stability^17^, allowing for prolonged reduction in pain behavior after a single local injection in various pain models^13,14,18^. Indeed, we observed weeks of analgesia after a single administration. In this study, our objective was to demonstrate the efficacy of intra-articular injected Na_V_1.8 channel decoy peptide, specifically designated as PY(A) peptide^13,16^, in alleviating osteoarthritis (OA) pain behavior. Additionally, we assessed the impact of the PY(A) peptide on OA-related bone pathophysiology.

## Results

### Expression of Magi-1 in Na_V_1.8^+^ joint afferents of human synovium

The intra-articular space is densely innervated with Na_V_1.8^+^ sensory neurons, which makes it an attractive target for local therapeutic approaches to treat OA pain^19^. Previously, we demonstrated co-localization of Magi-1 and Na_V_1.8 in the somata of human and mouse DRG neurons, as well as in skin peripheral nerve terminals^13,14^. Here, we found that Magi-1 was also expressed within Na_V_1.8^+^ nerve afferents innervating human synovium (Fig. 1). We also found Magi-1 expression in mouse knee subchondral bone (Supplementary Fig.1).

**Figure 1.**
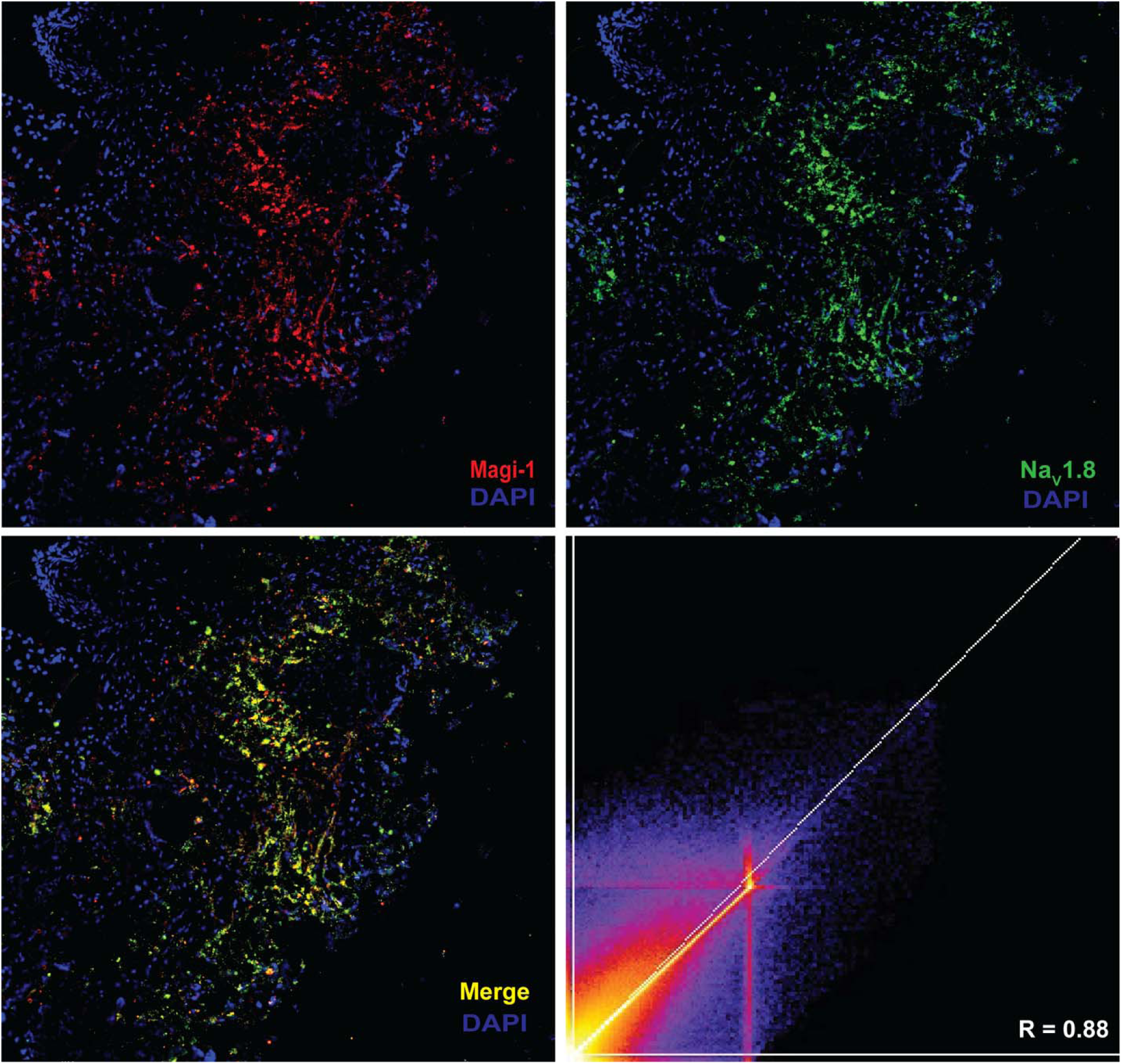
Scaffold protein Magi-1 is expressed in Na_V_1.8+ nociceptors innervating human synovium. Representative immunofluorescent image showing expression of Magi-1(red) and Na_V_1.8(green) in human synovium. Colocalization of Magi-1 and Na_V_1.8 is expressed as merge image (yellow). Scatter plot graph shows Pearson correlation coefficient (*R =* 0.88). Scale bar = 200 µm, 10X magnification.

### *In vivo* Magi-1 knockdown decreased OA pain-like behavior in mice

We used a sciatic nerve, Magi-1-targeted shRNA plasmid transfection knockdown method in mice ^13,14^ (Fig. 2a) to genetically support the role of Magi-1 in OA pain signaling. This knockdown approach resulted in a 70-75% reduction in Magi-1 protein (seven days post plasmid injection)^14^. We assessed whether Magi-1 deficiency affected pain behavior in the monoiodoacete (MIA) model of OA in mice^20^. MIA induces chondrocyte apoptosis by inhibiting glyceraldehyde-3-phosphatase dehydrogenase, disrupting glycolysis and cell survival^21^. This results in progressive articular cartilage degradation, release of pro-inflammatory cytokines, and abnormal subchondral bone remodeling^22^. In these experiments, mice were evaluated for pain behavior up to 20 days after MIA injection by dynamic weight bearing (DWB) and von Frey filament mechanical sensitivity assays (Fig. 2b). Mice received either Magi-1 or non-coding control shRNA plasmid injections on day 5 post MIA. Our results showed that Magi-1 knockdown mice improved weight bearing on the injured ipsilateral side compared to control mice on day 8 up to day 16 post MIA injection (Fig 2c). Additionally, significant increase in paw withdrawal threshold was observed in Magi-1 knockdown mice compared to control group (Fig 2d). Data was also segregated based on sex (Supplementary Fig. 2). These data validated the impact of Magi-1 deficiency on OA-mediated pain signaling.

**Figure 2.**
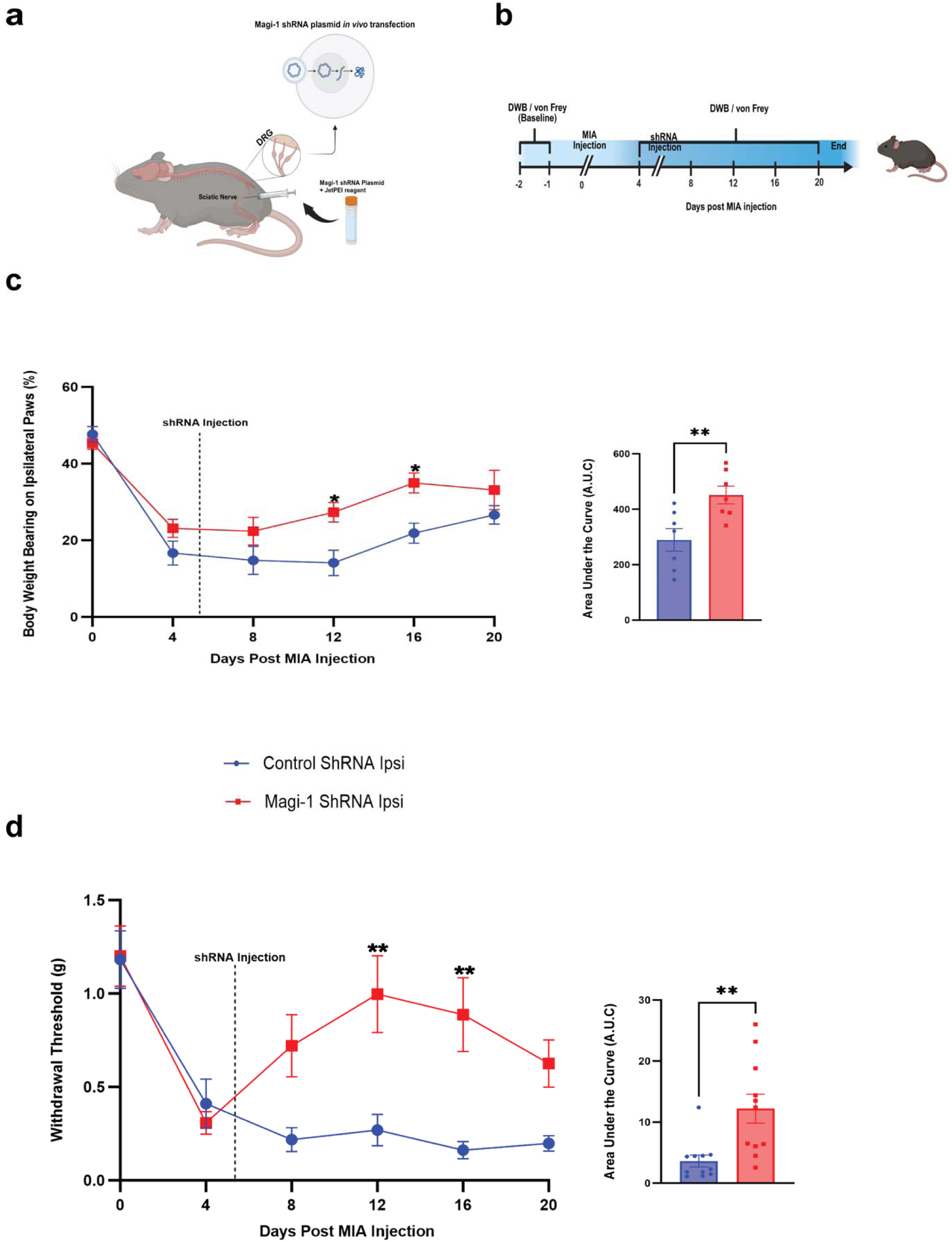
Magi-1 knockdown attenuates pain-like behavior in OA mice. **a** Schematic diagram showing the mouse model of *in vivo* Magi-1-targeted shRNA plasmid transfection. **b** Experimental timeline for pain behavior assessment. **c** Percent of weight borne on the ipsilateral paw of animals after MIA injection and Magi-1 or control shRNA plasmid. Pooled data for males and females is represented as cumulative mean ± S.E.M (*n* = 7 per group). Significance determined by repeated measures 2-way ANOVA with Bonferroni correction *p* < 0.05; \**p* < 0.01; \*\**p* < 0.001; *** (Magi-1 vs. Control). Total area under the curve (A.U.C) of animals assessed for ipsilateral weight bearing behavior. Significance determined by unpaired Student *t* test. **d** von Frey withdrawal threshold (g) of ipsilateral paw of animals represented as cumulative mean ± S.E.M (*n* = 11 per group). Significance determined by repeated measures 2-way ANOVA with Bonferroni correction *p* < 0.05; \**p* < 0.01; \*\**p* < 0.001; *** (Magi-1 vs. Control). Total area under the curve (A.U.C) for von Frey behavior. Significance determined by unpaired Student *t* test.

### Lipidated PY(A) peptide attenuated OA pain behavior in rats for weeks after a single intra-articular injection

Previously, we showed that Na_V_1.8-derived PY(A) lipidated peptide promoted the channel degradation in DRG neurons and inhibited neuropathic pain behavior in mice for up to 21 days after a single local administration^15^. Here, we evaluated whether an intra-articular injection of the PY(A) peptide into the inflamed knee joint could decrease pain behavior in OA rats. Animals were given a single intra-articular injection of PY(A) or scrambled peptide (200 µM, 50 µL) 5 days after OA induction and assessed for pain behavior up to 28 days post MIA (Fig. 3a). PY(A) peptide animals showed an increase in percentage weight bearing on the arthritic ipsilateral side compared to scrambled peptide animals for multiple weeks after peptide injection (Fig. 3b). Additionally, a compensatory decrease in weight bearing was simultaneously observed on the uninjured contralateral side of PY(A) animals compared to scrambled group (Fig. 3c). MIA-stimulated mechanical sensitivity was assessed using the von Frey assay. Animals injected with PY(A) peptide exhibited a significantly increased paw withdrawal threshold compared to scrambled animals 19 days after administration (Fig. 3c) with the effect wearing off by day 28 of the assay. Contralateral mechanical sensitivity was not different between groups (Supplementary Fig. 3). Minor differences were noted in the behavior assays performed after data was segregated by sex. (Supplementary Fig. 4). These data demonstrated that intra-articularly injected lipidated PY(A) peptide effectively reduced OA pain behavior for a prolonged duration.

**Figure 3.**
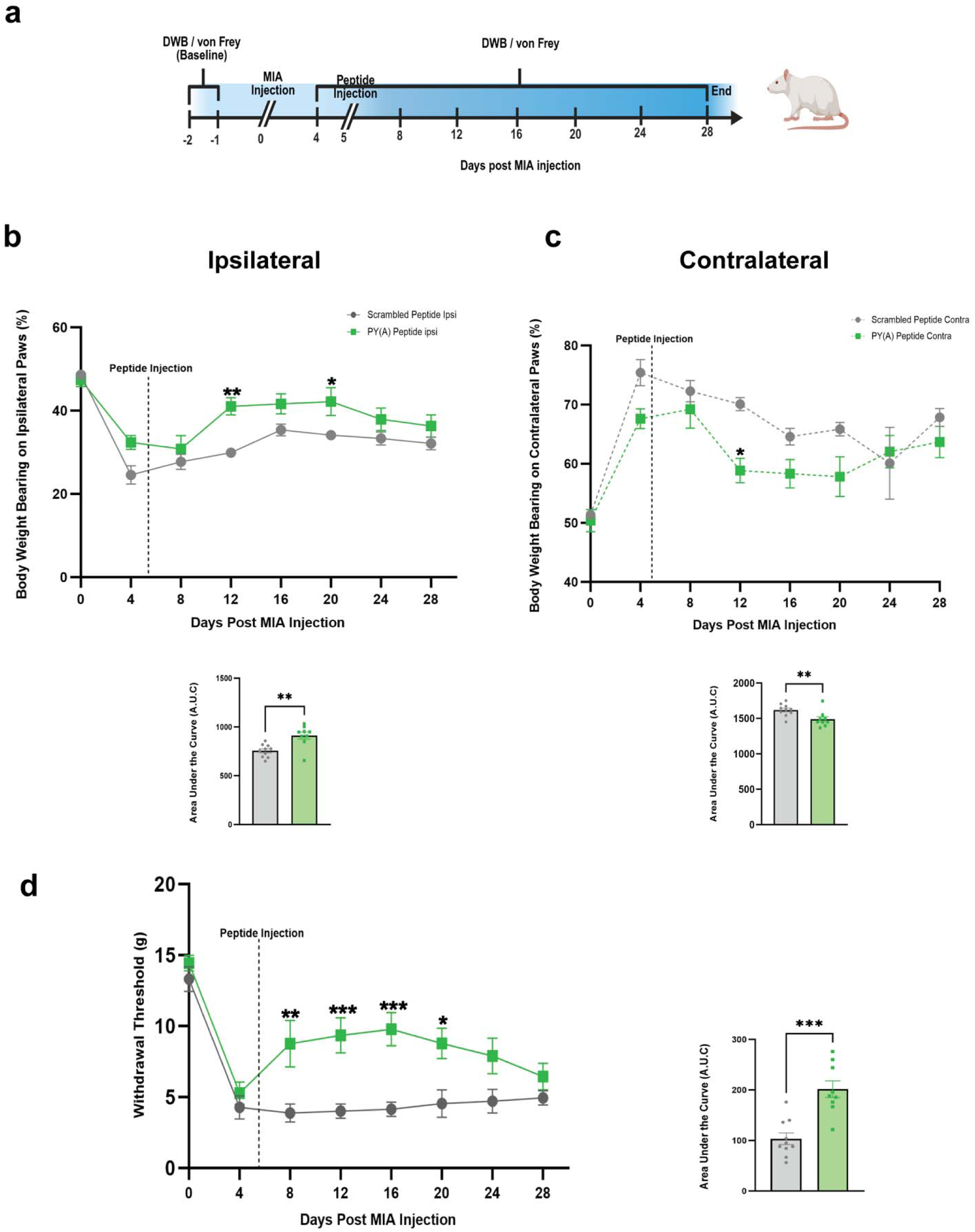
Local administration of PY(A) lipidated peptide reduced pain behavior in OA rats. **a** Experimental timeline for pain assessment. **b** Percent of weight borne on ipsilateral paw of MIA rats injected with PY(A) (*n* = 9) or scrambled peptide (*n* = 10). Pooled data for males and females is represented as cumulative mean ± S.E.M. Significance determined by repeated measures 2-way ANOVA with Bonferroni correction *p* < 0.05; \**p* < 0.01; \*\**p* < 0.001; *** (PY(A) vs. Scrambled). Total area under the curve (A.U.C) of animals assessed for ipsilateral weigh bearing behavior. Significance determined by unpaired Student *t* test. **c** Percent of weight borne on contralateral paw. Data represented as cumulative mean ± S.E.M. Significance determined by repeated measures 2-way ANOVA with Bonferroni correction *p* < 0.05; \**p* < 0.01; \*\**p* < 0.001; *** (PY(A) vs. Scrambled). Total area under the curve (A.U.C) of animals assessed for contralateral weight bearing behavior. Significance determined by unpaired Student *t* test. **d** von Frey withdrawal threshold (g) of ipsilateral paws of MIA animals injected with PY(A) (*n* = 9) or scrambled peptide (*n* = 10). Total area under the curve (A.U.C) for von Frey behavior. Significance determined by unpaired Student *t* test.

### Histological analyses of arthritic knees after intra-articular peptide injection

Knee joints from PY(A) and scrambled peptide treated animals were collected at the end of MIA assay (28 days after MIA injection). OA knee joints injected with the PY(A) peptide and scrambled peptide were evaluated for cartilage integrity by safranin-O staining and compared to healthy contralateral joints. Our results indicated both PY(A) peptide- and scrambled peptide-treated animals had substantial cartilage loss, typical of MIA treatment, when compared to healthy control contralateral knee joints. Furthermore, PY(A) peptide joints did not show any significant difference in cartilage degradation when compared to scrambled groups (Fig. 4a, 4b). These data verified that PY(A) peptide treated animals had arthritic knees, as evidenced by significant cartilage loss.

**Figure 4.**
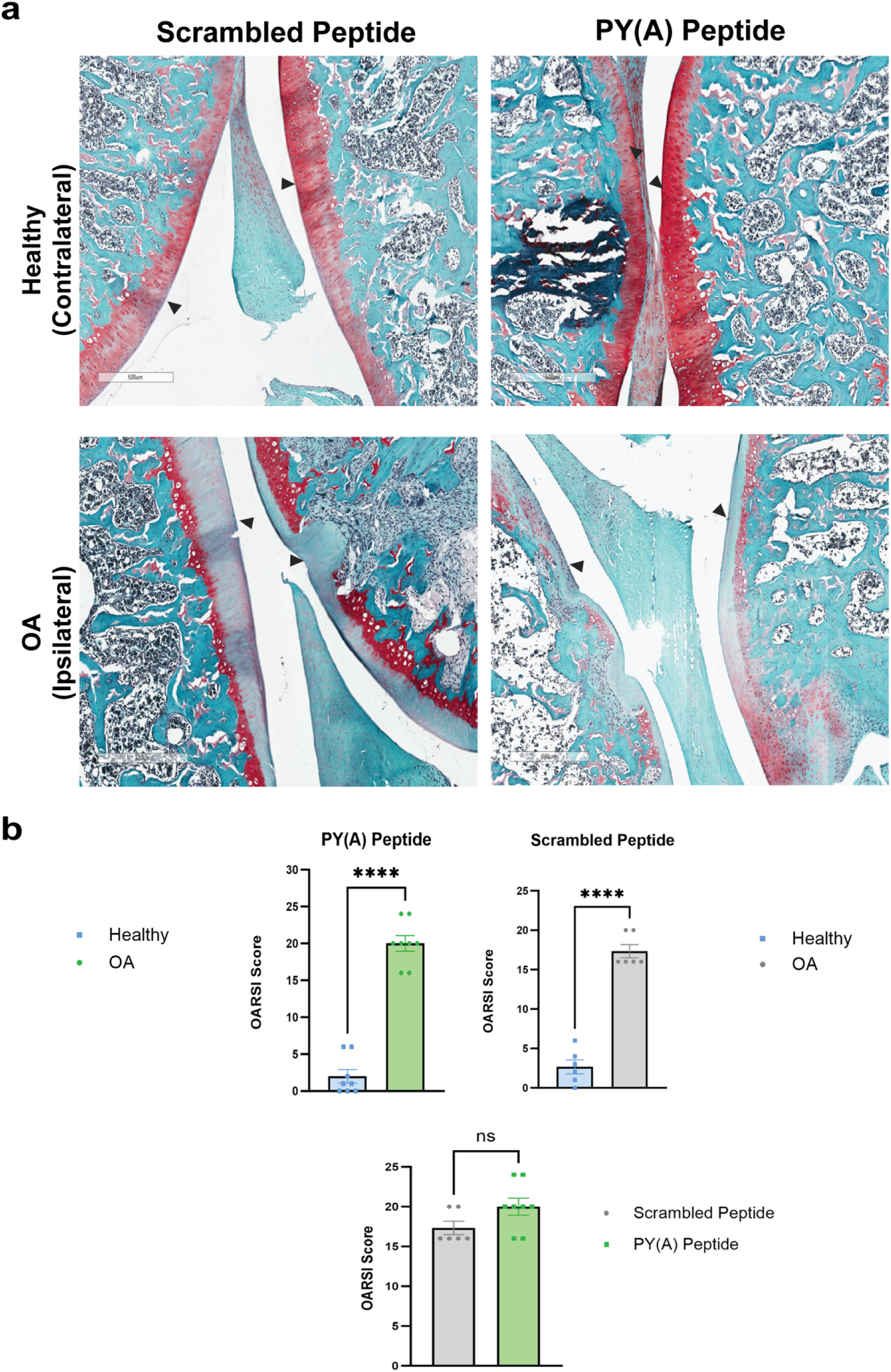
Lipidated PY(A) peptide does not accelerate OA pathology in rats. **a** Representative images of tibial cartilage (Safranin-O) 28 days post MIA injection. Samples received a single intra-articular injection of PY(A) (*n* = 8) or scrambled peptide (*n* = 6) at day 4 post MIA administration. Arrows indicate examined cartilage regions **b** Cartilage integrity was determined by the OARSI scoring system^25^ and represented as cumulative means S.E.M. Ipsilateral PY(A) and scrambled groups (OA) were compared to contralateral (healthy) groups. Significance was determined by unpaired Student *t* test. No significant differences were observed between PY(A) and scrambled groups.

### Lipidated PY(A) peptide reduced OA-induced abnormal subchondral bone remodeling

We further assessed joint integrity by performing a micro-computed tomography (MicroCT) analysis of arthritic rat knee joints injected with either PY(A) or scrambled peptide 28 days after MIA administration. Animals injected with scrambled peptide showed the typical MIA-induced erosion of subchondral bone^23^ with a significant reduction in bone volume fraction (BV/TV%) compared to healthy contralateral joints (Fig. 5a&c). To our surprise we observed no significant difference in BV/TV% between PY(A)-injected knee joints and healthy contralateral joints (Fig. 5b&d). Quantification of the MicroCT data showed a higher BV/TV% PY(A) peptide animals compared to scrambled peptide treated animals, (Fig. 5e). These data suggest that the sustained pain reduction provided by the PY(A) peptide delayed arthritic subchondral bone erosion.

**Figure 5.**
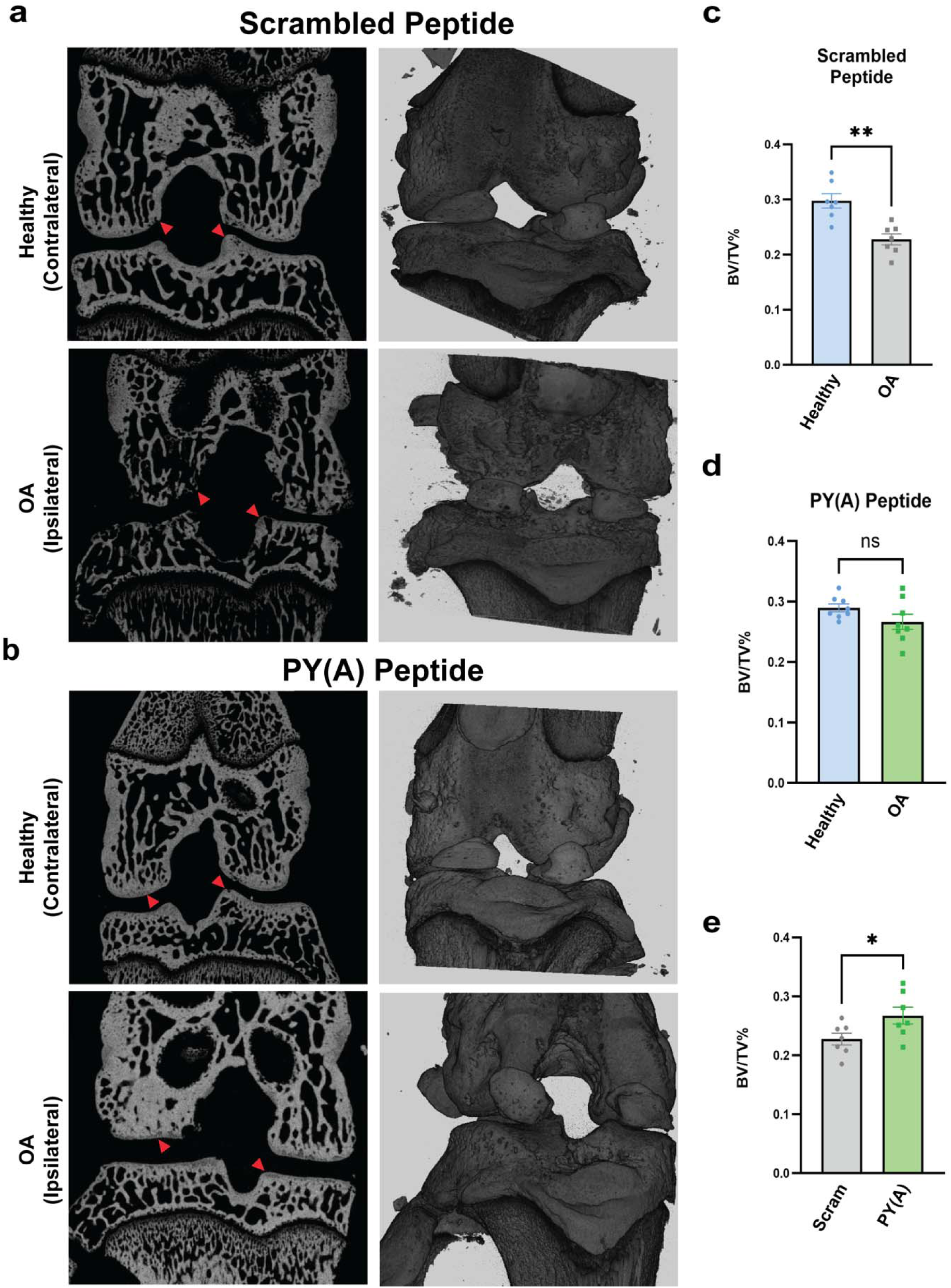
Pain inhibition by local lipidated PY(A) peptide reduces abnormal subchondral bone remodeling. **a** Representative micro-computed tomography (microCT) images of rat knee subchondral bone 28 days after MIA-induced OA. Animals received a single intra-articular injection of scrambled peptide (*n* = 7) on day 4 post MIA administration. **b** MicroCT images of knee subchondral bone of animals injected with PY(A) peptide (*n* = 7) on day 4 post MIA administration. **c** Scrambled group comparison of ipsilateral (OA) vs. contralateral (healthy) joints. **d** PY(A) group comparison of ipsilateral (OA) vs. contralateral (healthy) joints. **e** Comparison of PY(A) vs. scrambled groups. Bone volume fraction (BV/TV) analysis was performed using ImageJ plugin BoneJ and data was represented as cumulative means S.E.M. Significance was determined by unpaired Student *t* test. BV/TV significant differences were observed in scrambled vs. healthy groups, and PY(A) vs. scrambled groups. No significant differences were detected in PY(A) vs. healthy groups.

## Discussion

Previously, we demonstrated that small lipidated peptides, approximately 10-15 amino acids in length, readily penetrate peripheral nerve endings *in vivo*^18^. Furthermore, we showed that local injection of the lipidated PY(A) peptide effectively reduced neuropathic pain behavior for several weeks after a single dose^13^. Based on these findings, we anticipated that a single intra-articular injection of the PY(A) peptide into an arthritic joint would similarly attenuate osteoarthritis (OA) pain behavior for an extended period. This is precisely what we observed: robust attenuation of pain behavior for multiple weeks, indicating that small lipidated peptides targeting joint afferents hold promise as a novel therapeutic approach for OA pain. Unexpectedly, using microCT imaging, we also observed subchondral bone preservation during PY(A) peptide treatment. This implies that prolonged analgesia, in conjunction with the concurrent ability to utilize the arthritic joint, is essential for preventing bone loss.

Integrating supporting genetic evidence significantly boosts the success rate of clinical drug development^24^. Accordingly, we used an *in vivo* sciatic nerve gene knockdown approach in mice^13,14,18^ to test whether Magi-1 deficiency affected OA pain behavior. Magi-1 immunoreactivity was previously shown to be highest in the growth cones of cultured DRG neurons^25^ and in regenerating neurons growth cones contain some of the highest densities of sodium and calcium channels^26^. Our prior work also indicated that Magi-1 scaffolded Na_V_1.8 channels and protected channels from degradation in DRG neurons^14^. We found that Magi-1 deficiency resulted in increased weight bearing on the arthritic knee and decreased mechanical sensitivity (Fig 2) underpinning its role in OA pain signaling. RNAi-based therapeutics for cartilage diseases are experiencing numerous groundbreaking advancements^27^. Our evidence indicates that Magi-1 should be considered a potential siRNA target for the treatment of OA pain. Furthermore, our Magi-1 genetic data provided validations for our lipidated peptide pharmacological approach.

Intra-articular drug delivery offers numerous advantages, including direct access to the joint space. This enables the increased bioavailability of therapeutic agents at the affected site, while simultaneously reducing systemic exposure, potential side effects, and overall costs^28^. However, the therapeutic effectiveness of intra-articular injections remains severely limited due to rapid clearance of the drugs^28^. As we have previously shown, small lipidated peptides remain present in neuronal membrane for many weeks after administration^13^. A single intra-articular injection of the PY(A) peptide increased weight bearing on the injured arthritic side, decreased compensatory weight bearing on the uninjured contralateral side and decreased mechanical sensitivity in male and female rats for multiple weeks (Fig 3). It was necessary to use rats as the bigger animals allowed for more accurate intra-articular injection into the swollen knees. Even in clinical practice, ultrasound guidance notably improved injection accuracy in the target intra-articular joint space of knee joints, improving pain outcomes^29^. Furthermore, as previously demonstrated, the Magi-1 protection of Na_V_1.8 channels is conserved across multiple species, including humans^13,14^. In OA, nerve afferents located within the synovial membrane are the most significant source of pain due to sensitization caused by inflammatory responses in the joint environment, leading to activation of pain signals^30^. Our data indicates that intra-articular injection of PY(A) peptide readily accesses synovial afferents as evidenced by the observed reduction in OA pain behavior. It is noteworthy that synovial fluid contains albumin^31^ and lipidated peptides are predicted to bind to albumin^32^. It is uncertain whether the synovial albumin may have attenuated some of the observed effects or perhaps acted as a depot, contributing to the prolonged duration of action. Nonetheless, the PY(A) peptide exhibits potent disruption of Na_V_1.8 scaffolding, acting in the sub-micromolar range^13^. We administered a single dose of 200µM (∼18µg) of peptide, resulting in a substantial reduction in pain behavior that effectively overcame any potential confounding effects of synovial fluid albumin.

For rigor, after the completion of the MIA OA pain assay, we sacrificed animals and assessed cartilage levels through histological analyses and bone volume using microCT. Although MIA-induced pain behavior was confirmed prior to injection of peptide, this additional assessment ensured that all animals were experiencing MIA-induced cartilage loss, and that pain relief was not due to inadequate MIA action. Additionally, we sought to ensure that the PY(A) peptide was not aggravating joint degeneration. As shown in Figure 4, cartilage loss in scrambled and PY(A) peptide-treated animals was equivalent. Notably, the federal drug administration did not approve Pfizer’s tanezumab (anti-nerve growth factor (NGF) antibody) for the treatment of osteoarthritis pain for the primary reason that tanezumab caused accelerated joint destruction^33^ and many patients enrolled in clinical trials required joint replacement. On tanezumab, Miller et al., opined: “in light of the reported adverse events in clinical trials, it is clearly warranted that preclinical studies not only assess the effects on pain-related outcomes but also include assessment of the joint”^34^. For many of the preclinical anti-NGF antibody studies, joint histology studies were not performed^34^. Consequently, we undertook an additional step to also assess bone integrity through microCT analyses. Our primary objective was to investigate whether the PY(A) peptide was not exacerbating joint destruction. To our surprise, the PY(A) peptide exhibited a disease-modifying effect by reducing arthritic-induced bone erosion. Although the precise mechanism of action remains uncertain, it is plausible that the sustained analgesic effect which led to increased weight-bearing on the arthritic joint (Fig 3) contributed to the reduction in bone loss. Both clinical and preclinical studies have demonstrated that elevated physical activity and mechanical loading on the arthritic joint maintain bone remodeling homeostasis^35,36,36^. Indeed, the microCT findings regarding bone volume provided further support for the enduring analgesic efficacy of the PY(A) peptide, which implies that microCT analysis itself might also serve as a marker for analgesia for other preclinical OA-related drug studies. Considering these observations, our research demonstrates that small lipidated peptides targeting joint afferent excitability present a promising novel therapeutic approach for the management of OA pain.

## Materials and Methods

### Animals

8-10 weeks old C57BL/6 mice and 250-275 g Sprague Dawley rats were purchased from Envigo (Indianapolis, IN). All animals were housed separately on a 12 h light/dark cycle and provided with food and water ad libitum. All animal experiments were conducted during the light cycle. A seven-day acclimation period was allowed before performing any behavioral experiments. Sex differences between male and female animals were also considered for all behavioral experiments performed. All animal experiments performed were approved by The Institutional Animal Care and Use Committee (IACUC) at The State University of New York at Buffalo, and followed the guidelines established by the National Institute of Health (NIH). All experiments were performed using blinding to the experimental condition. Unblinding occurred after the analysis.

### Monoiodoacetate (MIA) Model of Osteoarthritis

Osteoarthritis was induced by intraarticular administration of MIA (Sigma-Aldrich, St Louis, MO)^18^. C57BL/6 mice were anesthetized with 2% isoflurane and placed on a heating pad in a supine position. Using a 30-gauge x ½ inch Hamilton syringe (Hamilton, Reno NV), mice were injected on the right knee intraarticularly with 0.8 mg MIA in 8 µL of PBS. A 28-gauge x ½ inch needle was used to inject Sprague Dawley rats with 2 mg MIA in 30 µL of PBS. To ensure the MIA stayed within the knee joint cavity, injection depth was kept at 2 mm for mice, and 5 mm for rats by using a polyethylene tube to prevent further needle penetration. The needle was maintained in the knee joint for at least 60s after injection to avoid backflow. All animals were observed until fully recovered and then placed back in the animal room.

### Magi-1-targeted *in vivo* sciatic nerve shRNA transfection

The *in vivo* shRNA transfection was performed according to protocols from previous studies^12^. C57BL/6 mice were anesthetized with 2% isoflurane and placed on a heating pad in a prone position. After the animal was shaved and disinfected, a posterior longitudinal incision was performed at the lumbar spine region. Using sterile toothpicks, muscles around the lumbar spine were separated and sciatic nerve was exposed. 3 µL of Magi-1 shRNA plasmid or control were injected into the sciatic nerve using a Hamilton syringe with a 32-gauge blunt needle. Magi-1 shRNA and control were purchased from Santa Cruz Biotechnology (Santa Cruz, CA, USA). After the injection, the incision was closed with wound clips and animals were observed until full recovery.

### Lipidated Na_V_1.8 decoy peptide

N-terminal myristoylated peptides were designed based on the Na_V_1.8 PY motif in the C-terminal with a threonine to alanine substitution (SATSFPPSYDSV<u>A</u>RG) (PY(A) peptide)^13,14^. The scrambled control peptide (SDRPYTSYSFSAPGT) was similarly myristoylated at the N-terminal. All peptides were synthesized by GenScript (Piscataway, NJ). 1 mg of lyophilized peptides were dissolved in DMSO to make a stock working solution. A final concentration of 200 µM lipidated peptide (DMSO <0.05%) was achieved by adding PBS. 100 µL aliquots were stored at −20°C until needed. Male and female rats were evaluated behaviorally 4 days post MIA injection to confirm knee pain sensitivity. Next day, animals were placed in the surgery room and anesthetized with 2% isoflurane. Once ipsilateral knee joint was sterilized, animals were injected intraarticularly with 50 µL of lipidated peptidomimetic (200 µM) using a 28-gauge x ½ inch Hamilton syringe. The needle was kept in the knee joint for at least 60 s to prevent peptide leakage. All animals were observed until full recovery and then placed back in the animal room.

### Pain Behavior Assays

#### Mechanical Sensitivity

Animals were enclosed on top of a testing grid surface (Ugo Basile) and allowed 30 minutes to acclimate to the testing chamber. Using Touch Test Sensory Probes (Stoeltings, Wood Dale, IL), a force was applied to the ipsilateral and contralateral plantar surface of the hind paw of the animals. Filaments were applied according to the Simplified Up-Down method (SUDO)^24^. An initial filament size was presented to the animal, and withdrawal response was recorded as positive or negative. A positive response resulted in a smaller filament size application, while a negative response resulted in a larger filament size application. A total of five filament presentations were used per animal with an interval of 5 minutes per filament. The last filament response was used to calculate withdrawal threshold based on equations described in previous protocol^24^.

#### Dynamic Weight Bearing

After 30 minutes of acclimation in the testing room, animals were weighed and placed into a Dynamic Weight Bearing (DWB) apparatus (BIOSEB, France). The DWB chamber was connected to a sensor that recorded weight exerted by the paws of the animals, and a video camera that monitored animal activity during the testing period. A red plexiglass sheet was placed around the DWB chamber to create a uniform environment for the animal. A 180 s latency period was allowed for animals to explore the chamber, and this was followed by a 300 s acquisition period. At the end of the recording, acquisition data was manually validated for 90 seconds randomly to accurately identify the weights coming from each paw.

### Histology

Rat knee joints were collected 28 days after MIA injection and fixed with 4% paraformaldehyde (PFA) for 48 h at 4°C. Sample decalcification was performed with 10% EDTA for 14 days. Decalcified samples were washed with PBS and embedded in paraffin for sectioning. 4 µm-thick sagittal sections were placed on slides and stained with safranin-O. Section images were captured at 20X magnification with Aperio CS2 slide scanner (Leica Biosystems, Germany). Tibial cartilage was scored according to the OARSI scoring system guidelines^25^.

### Micro-computed Tomography (microCT)

Knee joints were collected from rats 28 days after MIA injection. Samples were fixed with 4% PFA for 48 h and then washed with PBS. 70% ethanol was added to each sample and then placed in a ScanCo µCT100 scanner for analysis (Scanco Medical, Brüttisellen, Switzerland). Scanner parameters were configured as 70kVp for X-ray voltage, while image resolution was 10 µm. For subchondral bone analysis, reconstructed DICOM files were imported to the ImageJ software. Tibial subchondral bone was chosen as the region of interest for remodeling analysis. Changes in subchondral bone remodeling were analyzed based on bone volume fraction (BV/TV) using ImageJ plugin BoneJ (NIH).

### Immunohistochemistry

Human synovium tissue was donated by a 73-year-old female patient who underwent knee joint replacement surgery at Buffalo General Medical Center (Buffalo, NY). Tissue donation was consistent with institutional review board (IRB) guidelines according to protocol as approved: Orthopedic Tissue Procurement University at Buffalo IRB STUDY1729. No other identifying data was collected. Mouse knee joints were collected following a transcardial perfusion and fixation previously described in a protocol. Samples were incubated with 4% PFA for 24 h at 4°C. Knee joint decalcification was achieved by incubation in 10% EDTA for 14 days at 4°C. Then, fixed samples were frozen in Tissue-Tek OCT freezing medium (Sakura Finetek, Torrance, CA). Samples were sectioned at 20 µm (human synovium) and 30 µm (mouse knee joint) and placed on microscope slides (Fisherbrand, Pittsburgh, PA). Sections were washed 3 times with wash buffer (0.4% Triton X-100 in PBS), then incubated overnight in blocking solution (10% normal goat serum, 3% bovine serum albumin, and 0.4% Triton X-100 in PBS). The next day, slides were incubated in primary antibodies overnight at 4°C (Monoclonal Mouse anti-Magi-1, Novus Biologicals 1:500, Rabbit Polyclonal anti-Na_V_1.8, Abcam 1:500, Rabbit Polyclonal anti-Peripherin Abcam, 1:500). The following day, slides were incubated in secondary antibodies overnight at 4°C (Donkey anti-mouse 546, Invitrogen 1:1000, Donkey anti-rabbit 647, Abcam 1:1000). Afterwards, slides were washed twice with wash buffer and mounted using Vectashield antifade mounting medium with DAPI (VectorLabs, Newark, CA). All images were acquired using a Leica DMi8 inverted fluorescent microscope (Leica Biosystems, Germany), and analyzed using the LAS X imaging software. Pearson correlation coefficient was analyzed using ImageJ.

### Statistics

All statistical analysis was performed using GraphPad Prism (GraphPad, La Jolla, CA). Statistical significance was evaluated using a *p-*value <0.05, and data was shown as means ± S.E.M. Comparison between two groups was assessed by unpaired Student *t* test, while 2-way ANOVA with Bonferroni correction was used for multiple comparison analysis.

## Data availability

Source for data generated in this study will be provided upon request.

## Acknowledgments

This work was supported by the National Institute of Health Heal Initiative grant NS113991 and NS128543. The authors would like to thank Dr. Imtiaz Mohammed of the University at Buffalo Histology Core facility for his assistance with knee joint safranin-O staining. We would also like to thank Dr. Andrew McCall for his assistance with MicroCT scanning at the Optical Imaging and Analysis facility. Included illustrations were created using BioRender.com.

## Conflict of interests

AB is a co-founder of Channavix Therapeutics, LLC and Mimetic Medicines, INC. A Patent Cooperation Treaty (PCT) application (serial number PCT/US2018/65545) was filed on the use of lipidated peptidomimetics targeting Na_V_1.8 to induce local analgesia and to treat pain by the University at Buffalo. RR is interning at Mimetic Medicines, INC. All other authors declare no competing interests.

## Author Contributions

AB and RR conceived the idea for the project and co-wrote the manuscript. RR performed all the behavioral, histology, knee imaging and immunohistochemistry experiments, analyzed all the data, and generated the manuscript. JBK isolated human synovial tissue.

## References

1 Hunter, D. J., Schofield, D. & Callander, E. The individual and socioeconomic impact of osteoarthritis. Nat Rev Rheumatol 10, 437–441 (2014).

2 Steinmetz, J. D. et al. Global, regional, and national burden of osteoarthritis, 1990&#x2013;2020 and projections to 2050: a systematic analysis for the Global Burden of Disease Study 2021. The Lancet Rheumatology 5, e508–e522 (2023).

3 Fu, K., Robbins, S. R. & McDougall, J. J. Osteoarthritis: the genesis of pain. Rheumatology 57, iv43–iv50 (2017).

4 Katz, J. N., Arant, K. R. & Loeser, R. F. Diagnosis and Treatment of Hip and Knee Osteoarthritis: A Review. JAMA 325, 568–578 (2021).

5 Billesberger, L. M., Fisher, K. M., Qadri, Y. J. & Boortz-Marx, R. L. Procedural Treatments for Knee Osteoarthritis: A Review of Current Injectable Therapies. Pain Res Manag 2020, 3873098 (2020).

6 McAlindon, T. E. et al. Effect of Intra-articular Triamcinolone vs Saline on Knee Cartilage Volume and Pain in Patients With Knee Osteoarthritis: A Randomized Clinical Trial. JAMA 317, 1967–1975 (2017).

7 Schuelert, N. & McDougall, J. J. Involvement of Nav 1.8 sodium ion channels in the transduction of mechanical pain in a rodent model of osteoarthritis. Arthritis Res Ther 14, R5 (2012).

8 Jones, J. et al. Selective Inhibition of Na(V)1.8 with VX-548 for Acute Pain. N Engl J Med 389, 393–405 (2023).

9 Wallace, M. S. Trials for Managing Acute Pain - A Clinically Meaningful Small Effect Size? N Engl J Med 389, 464–465 (2023).

10 Jo, S., Zhang, H. B. & Bean, B. P. Use-Dependent Relief of Inhibition of Nav1.8 Channels by A-887826. Mol Pharmacol 103, 221–229 (2023).

11 Vaelli, P. et al. State-Dependent Inhibition of Nav1.8 Sodium Channels by VX-150 and VX-548. Mol Pharmacol 106, 298–308 (2024).

12 Hang Kong, A. Y., Tan, H. S. & Habib, A. S. VX-548 in the treatment of acute pain. Pain Manag 14, 477–486 (2024).

13 Martin, M. K. et al. Pharmacologically enabling the degradation of NaV1.8 channels to reduce neuropathic pain. PAIN, 10.1097/j.pain.0000000000003470 (2024).

14 Pryce, K. D. et al. Magi-1 scaffolds Na(V)1.8 and Slack K(Na) channels in dorsal root ganglion neurons regulating excitability and pain. Faseb j 33, 7315–7330 (2019).

15 Ingham, R. J., Gish, G. & Pawson, T. The Nedd4 family of E3 ubiquitin ligases: functional diversity within a common modular architecture. Oncogene 23, 1972–1984 (2004).

16 Gomez, K., Calderon-Rivera, A. & Khanna, R. Pain’s puzzle pieces: MAGI-1, NaV1.8, degradation, analgesia. Pain (2024).

17 Menacho-Melgar, R., Decker, J. S., Hennigan, J. N. & Lynch, M. D. A review of lipidation in the development of advanced protein and peptide therapeutics. J Control Release 295, 1–12 (2019).

18 Powell, R. et al. Inhibiting endocytosis in CGRP(+) nociceptors attenuates inflammatory pain-like behavior. Nat Commun 12, 5812 (2021).

19 Strickland, I. T. et al. Changes in the expression of NaV1.7, NaV1.8 and NaV1.9 in a distinct population of dorsal root ganglia innervating the rat knee joint in a model of chronic inflammatory joint pain. Eur J Pain 12, 564–572 (2008).

20 Pitcher, T., Sousa-Valente, J. & Malcangio, M. The Monoiodoacetate Model of Osteoarthritis Pain in the Mouse. J Vis Exp (2016).

21 de Sousa Valente, J. The Pharmacology of Pain Associated With the Monoiodoacetate Model of Osteoarthritis. Front Pharmacol 10, 974 (2019).

22 Bao, Z. et al. Monosodium iodoacetate-induced subchondral bone microstructure and inflammatory changes in an animal model of osteoarthritis. Open Life Sci 17, 781–793 (2022).

23 Xie, L. et al. Quantitative imaging of cartilage and bone morphology, reactive oxygen species, and vascularization in a rodent model of osteoarthritis. Arthritis Rheum 64, 1899–1908 (2012).

24 Nelson, M. R. et al. The support of human genetic evidence for approved drug indications. Nat Genet 47, 856–860 (2015).

25 Ito, H. et al. Biochemical and morphological characterization of MAGI-1 in neuronal tissue. J Neurosci Res 90, 1776–1781 (2012).

26 Chen, D., Yu, S. P. & Wei, L. Ion channels in regulation of neuronal regenerative activities. Transl Stroke Res 5, 156–162 (2014).

27 Zhou, L., Rubin, L. E., Liu, C. & Chen, Y. Short interfering RNA (siRNA)-Based Therapeutics for Cartilage Diseases. Regen Eng Transl Med 7, 283–290 (2020).

28 Rai, M. F. & Pham, C. T. Intra-articular drug delivery systems for joint diseases. Curr Opin Pharmacol 40, 67–73 (2018).

29 Berkoff, D. J., Miller, L. E. & Block, J. E. Clinical utility of ultrasound guidance for intra-articular knee injections: a review. Clin Interv Aging 7, 89–95 (2012).

30 Nanus, D. E. et al. Synovial tissue from sites of joint pain in knee osteoarthritis patients exhibits a differential phenotype with distinct fibroblast subsets. EBioMedicine 72, 103618 (2021).

31 Bennike, T. et al. A normative study of the synovial fluid proteome from healthy porcine knee joints. J Proteome Res 13, 4377–4387 (2014).

32 Wang, Y. et al. The molecular basis for the prolonged blood circulation of lipidated incretin peptides: Peptide oligomerization or binding to serum albumin? J Control Release 241, 25–33 (2016).

33 Hochberg, M. C. Serious joint-related adverse events in randomized controlled trials of anti-nerve growth factor monoclonal antibodies. Osteoarthritis Cartilage 23 Suppl 1, S18–21 (2015).

34 Miller, R. E., Block, J. A. & Malfait, A. M. Nerve growth factor blockade for the management of osteoarthritis pain: what can we learn from clinical trials and preclinical models? Curr Opin Rheumatol 29, 110–118 (2017).

35 Liphardt, A. M. et al. Changes in mechanical loading affect arthritis-induced bone loss in mice. Bone 131, 115149 (2020).

36 Rausch Osthoff, A. K., et al. 2018 EULAR recommendations for physical activity in people with inflammatory arthritis and osteoarthritis. Ann Rheum Dis 77, 1251–1260 (2018).

