## Supplemental Figure 1 for "A small lipidated peptide targeting Na_V_1.8 channels attenuates osteoarthritic pain behavior and prevents bone erosion"

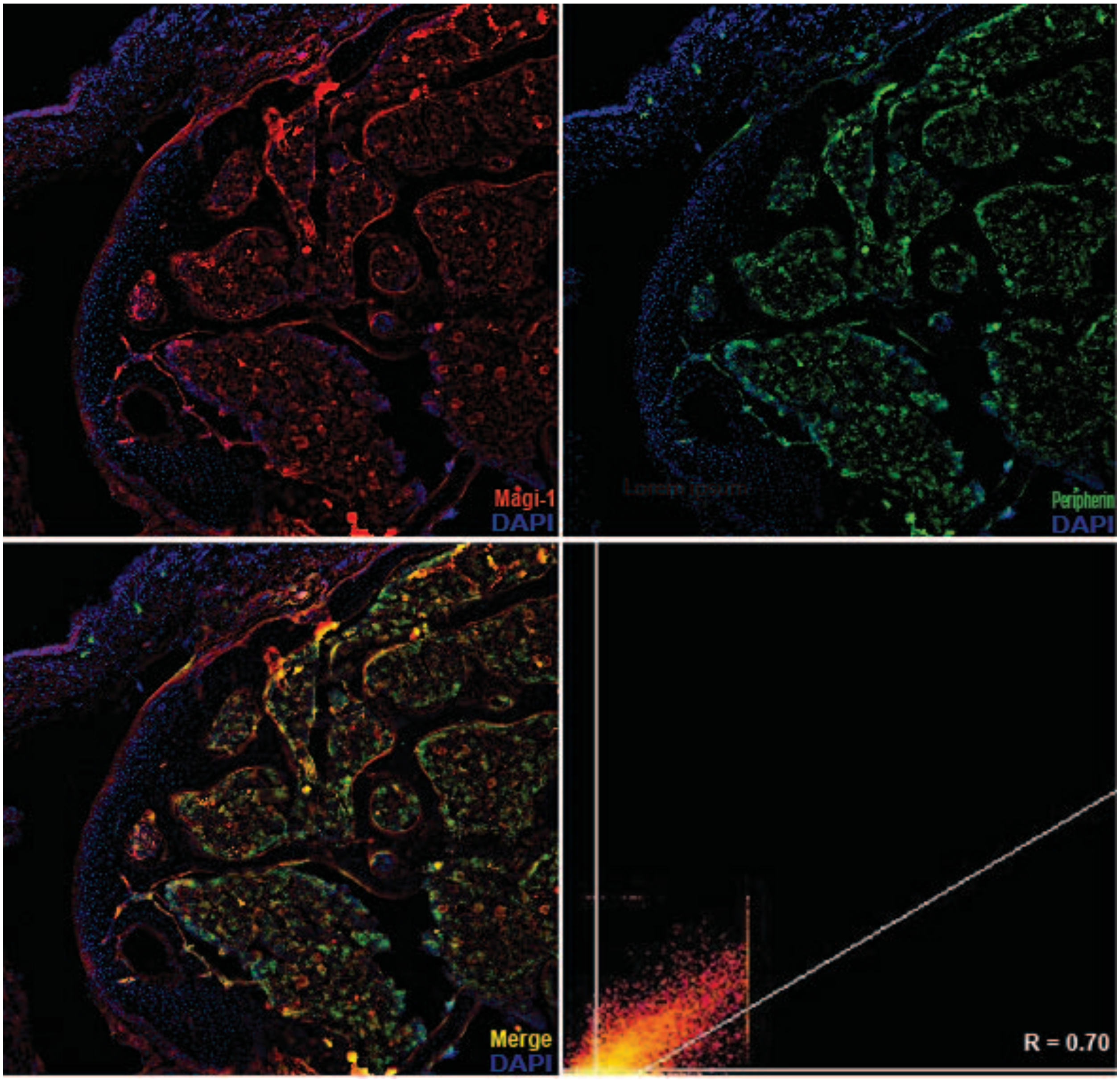


**Supplementary Figure 1. Representative immunofluorescent image of Magi-1 and peripherin in mouse knee subchondral bone nociceptors**. Colocalization of Magi-1 (red) and peripherin (green) is shown as merge image (yellow). Scatter plot graph shows Pearson correlation coefficient (*R* = 0.70). Scale bar = 200 μm, 10X magnification.
