## Supplemental Figure 2 for "A small lipidated peptide targeting Na_V_1.8 channels attenuates osteoarthritic pain behavior and prevents bone erosion"

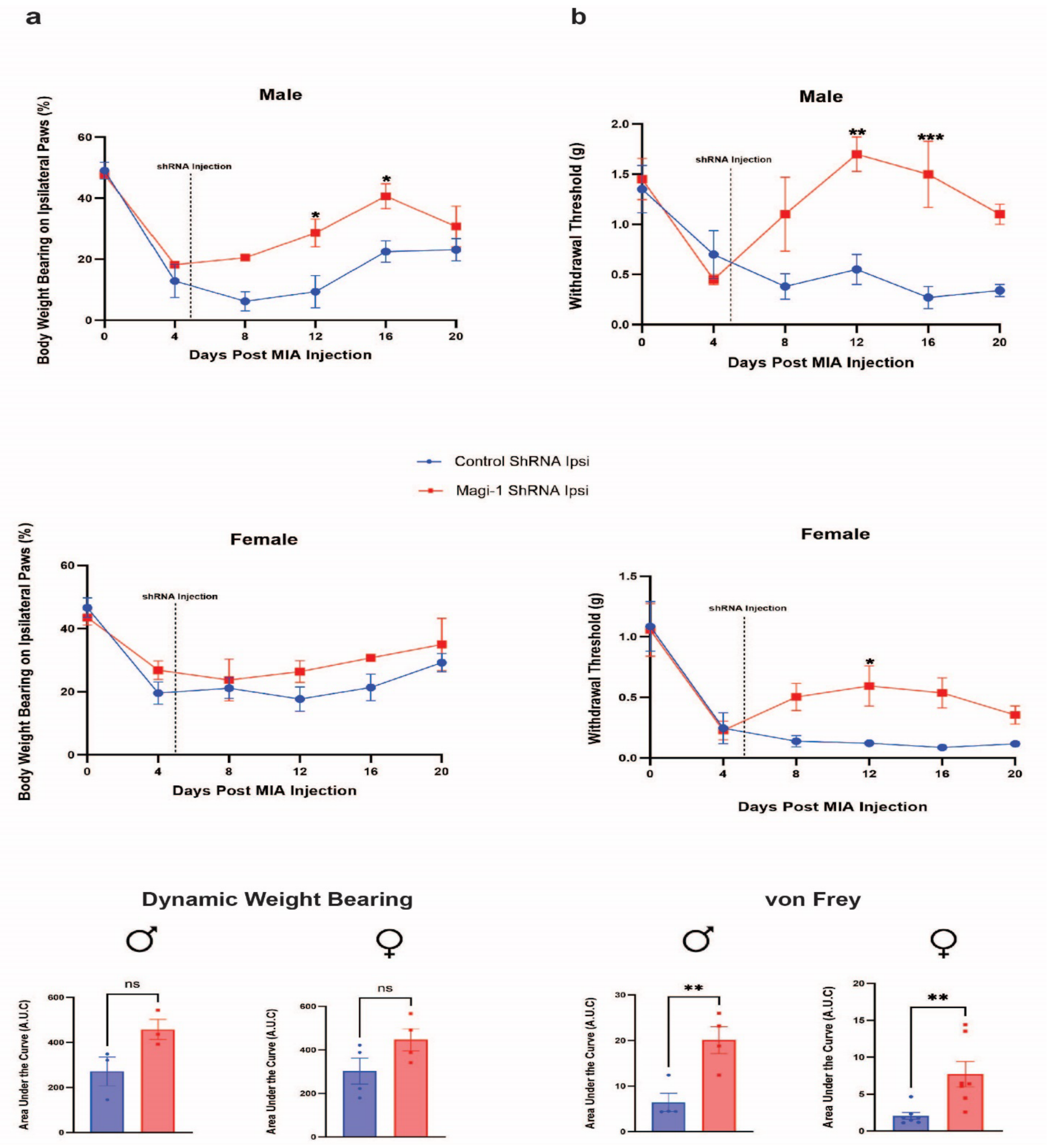


**Supplementary Figure 2. Sex segregation of pain-like behavior in Magi-1 knockdown OA mice. a** Percent of weight borne on the ipsilateral paw of male ($n$ = 3 per group) and female ($n$ = 4 per group) mice after MIA injection and Magi-1 or control shRNA plasmid. Data represented as cumulative mean ± S.E.M. Significance determined by repeated measures 2-way ANOVA with Bonferroni correction $p < 0.05$; *$p < 0.01$; **$p < 0.001$; *** (Magi-1 vs. Control). Total area under the curve (A.U.C) for weight bearing behavior assessment (male and female). Significance determined by unpaired Student *t* test. **b** von Frey withdrawal threshold (g) of ipsilateral paw of male ($n = 4$ per group) and female ($n = 7$ per group) mice represented as cumulative mean ± S.E.M. Significance determined by repeated measures 2-way ANOVA with Bonferroni correction $p < 0.05$; *$p < 0.01$; **$p < 0.001$; *** (Magi-1 vs. Control). Total area under the curve (A.U.C) for von Frey behavior (male and female). Significance determined by unpaired Student *t* test.
