## Supplemental Figure 3 for "A small lipidated peptide targeting Na_V_1.8 channels attenuates osteoarthritic pain behavior and prevents bone erosion"

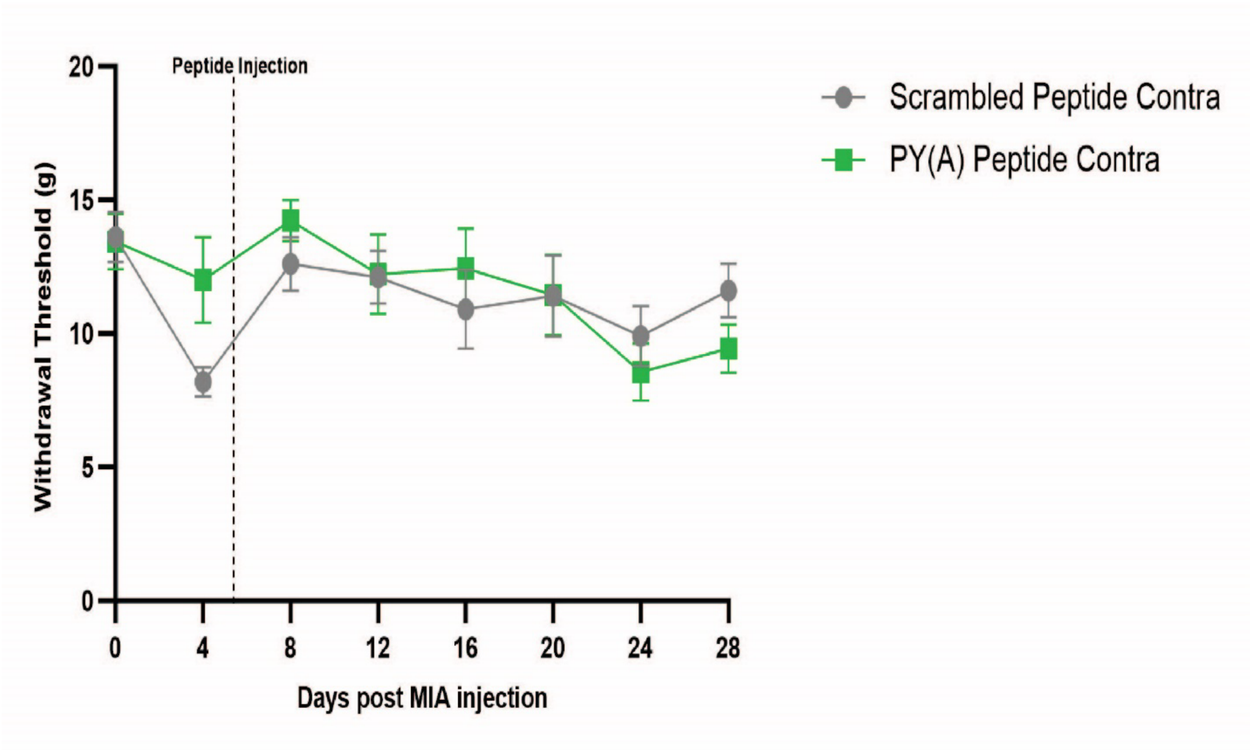


**Supplementary Figure 3. Contralateral pain behavior response in OA rats injected with PY(A) peptide.** von Frey withdrawal threshold (g) of contralateral paws of MIA animals injected with PY(A) ($n$ = 9) or scrambled peptide ($n$ = 10). Pooled data represented as cumulative mean ± S.E.M. Significance determined by repeated measures 2-way ANOVA with Bonferroni correction $p < 0.05$; *$p < 0.01$; **$p < 0.001$; *** (PY(A) vs. Scrambled). No significance was observed between groups.
