## Supplemental Figure 4 for "A small lipidated peptide targeting Na_V_1.8 channels attenuates osteoarthritic pain behavior and prevents bone erosion"

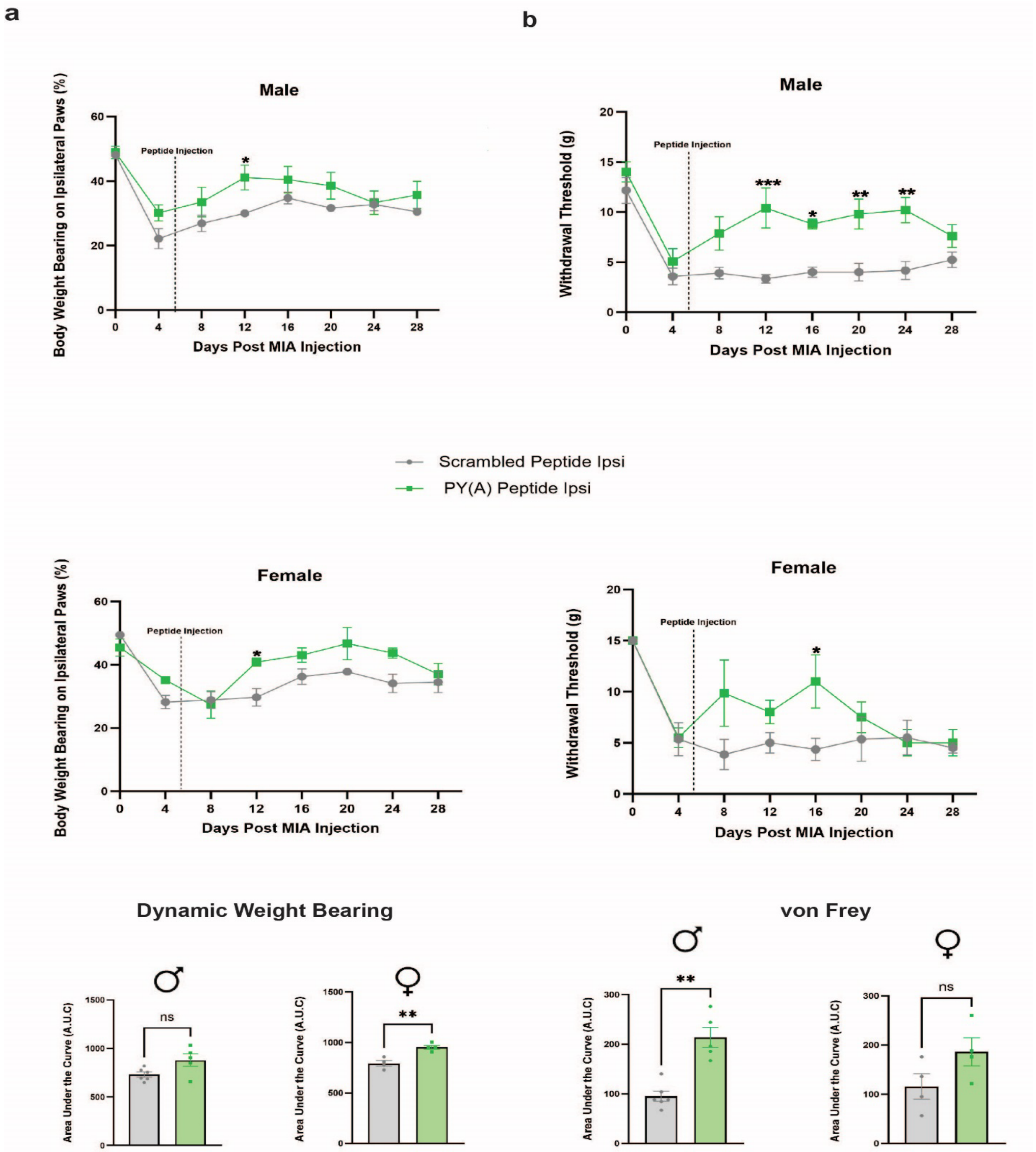


**Supplementary Figure 4. Sex segregation of pain-like behavior in OA rats injected with PY(A) peptide. a** Percent of weight borne on ipsilateral paw of OA rats injected with **PY(A)** (*n* = 5 (male), *n* = 4 (female)) or **scrambled** (*n* = 6 (male), *n* = 3 (female)) peptide.

Data represented as cumulative mean ± S.E.M. Significance determined by repeated measures 2-way ANOVA with Bonferroni correction $p < 0.05$; *$p < 0.01$; **$p < 0.001$; *** (PY(A) vs. Scrambled). Total area under the curve (A.U.C) of male and female animals for weight bearing behavior. Significance determined by unpaired Student *t* test. **b** von Frey withdrawal threshold (g) of ipsilateral paws of OA animals injected with **PY(A)** ($n = 5$ (male), $n = 4$ (female)) or **scrambled** ($n = 6$ (male), $n = 4$ (female)) peptide. Data represented as cumulative mean ± S.E.M. Significance determined by repeated measures 2-way ANOVA with Bonferroni correction $p < 0.05$; *$p < 0.01$; **$p < 0.001$; *** (PY(A) vs. Scrambled). Total area under the curve (A.U.C) of male and female animals for von Frey behavior. Significance determined by unpaired Student *t* test.
